# Fermented polyherbal formulation restored ricinoleic acid induced diarrhea in Sprague Dawley rats and exhibited *in vitro* antibacterial effect on multiple antibiotics resistant gastric pathogens

**DOI:** 10.1101/2023.11.22.568303

**Authors:** Subhanil Chakraborty, Babli Roy, Subhajit Sen, Santi M. Mandal, Ranadhir Chakraborty

## Abstract

The involvement of multiple antibiotics resistant gastric pathogens in diarrhea aggravates the disease condition uncontrollably. Current study was aimed at finding and developing a suitable formulation utilizing multiple natural components from known plant sources to augment current therapeutic outcomes. Hydro-ethanolic extraction method was applied through boiling and fermentation, on ancient observation based efficacious plant parts for developing antidiarrheal polyherbal formulation AP-01. Animal study model of diarrhea was used to evaluate safety and efficacy of the formulation. The formulation was tested *in vitro* on four different multiple antibiotics resistant gastric pathogens collected from national repository. The formulation depicted no cytotoxicity on normal gut cells and was efficacious at 10 ml/Kg single dose in relieving symptoms of diarrhea by 79.71%, in comparison with standard drug showing reduction of symptoms by 83.01%. AP-01 exhibited delaying the gastric motility. Symptoms of diarrhea ceased to occur within 321 minutes with AP-01, where the standard drug took 308 minutes. AP-01 was found successful at a viable dosage regimen of 75 to 100 microliter per ml, in inhibiting growth of different pathogens from Enterobacteriaceae family possessing resistance against several classes of antibiotics, in culture media. Chemical analysis revealed different alkaloids, flavonoids and polyphenols those probably work in unison through multiplex modes of action to arrest diarrhea and inhibit pathogens at the same time. These promising results shown by AP-01 should definitely evoke an effort to dive deep into research and development for better therapeutic formulations for infectious diarrhea by harvesting the arsenal of nature.

## Introduction

Gastrointestinal diseases, such as diarrhea, can easily overwhelm the public health system in developing countries. They spread quickly through food or drinking water and can lead to epidemics that cause significant loss of life even today.

Castor oil-induced diarrhea is often used in pre-clinical research to test drugs that control diarrhea because it provides a standardized and controllable method to induce diarrhea in laboratory animals, typically rodents [1–2]. Castor oil contains ricinoleic acid that irritates the intestines and stimulates bowel movements, leading to diarrhea[3]. Castor-oil-induced diarrhea typically has a predictable onset time after administration, allowing researchers to study the drug’s onset of action and its ability to relieve diarrhea symptoms. It is a cost-effective and efficient method of screening potential anti-diarrheal medications before moving on to more complex and expensive clinical trials. The ability to test drugs in a controlled laboratory setting using the castor oil-induced diarrhea model is a crucial step in the drug development pipeline [4-[5].

Infectious diarrhea, mainly due to pathogenic bacteria, is a significant public health problem worldwide. Major enteric bacterial pathogens responsible for causing diarrhea among human infants and young animals include several species of genera such as *Escherichia, Vibrio, Shigella, Salmonella, Campylobacter, Arcobacter, Yersinia, and Listeria*. Even within the genus *Escherichia, diarrheagenic Escherichia coli* (DEC) pathotypes, particularly *enteropathogenic E. coli* (EPEC), *enterohemorrhagic E. coli* (EHEC), *enterotoxigenic E. coli* (ETEC), *enteroaggregative E. Coli* (EAEC) and *entero-invasive E. coli* (EIEC) are frequently associated with diarrheal cases [7]. In developing nations, including India, there is a big hindrance to the availability of all the standard cultures for diagnosis and proper medication to treat the ailment. Even though economic development and progress in healthcare delivery are expected to catalyze substantial improvements in infectious disease-related morbidity and mortality by the year 2020, it is predicted that diarrhea will remain a leading health problem. The main cause of death from diarrhea is dehydration, which results from loss of electrolytes in diarrheal stools[8]. During the past decade, there have been some major improvements in treating infectious diarrhea. Oral rehydration therapy has contributed greatly to the reduction of diarrheal mortality rates in children and older people. However, the diarrheal attack rate has remained unchanged, and this treatment often fails in the high stool output state. Symptomatic therapy with antimotility agents is restricted to non-dehydrated patients without features of systemic infection. These are not indicated in infants and patients with febrile bloody diarrhea. Moreover, there is an increasing threat of drug resistance, side effects of antibiotic treatment, and supra infection when normal flora is eradicated with antimicrobial agents. Vaccines have been considered as the most feasible approach to diarrheal management. Various attempts have been made to develop vaccines against diarrhea-causing organisms [9–11]. However, the response to such vaccines in developing countries, particularly for toddlers, has not been encouraging[12]. Considerable technical barriers need to be overcome before continuing clinical evaluation of prospective vaccine candidates. Thus, an important niche exists for developing cost-effective alternative approaches, and medicinal plants may serve to fulfill this niche.

Various medicinal plant preparations have been identified for treating diarrhea, but their mechanism for eliminating organisms that cause diarrheal diseases is poorly comprehended. While data are available on the effect of different plants on intestinal motility in experimental models and antibacterial action [13–14], there is a lack of information on the efficacy of more than one medicinal plant working synergistically as a polyherbal formulation on various aspects of diarrheal pathogenicity[15]. The current study is aimed to find a suitable arsenal from nature with the help of ancient knowledge of Ayurveda and to introspect into the safety and efficacy of such preparation through in vivo and in vitro studies and to shed some light on the chemical constituents attributed to such actions.

## Materials and methods

### Plant parts for preparing fermented polyherbal formulation

Fermented Polyherbal formulation (AP-01) was prepared in the laboratory by boiling dried and cleaned stem barks of *Holarrhena antidysenterica* & *Gmelina arborea* mixed with large-sized dry resins of *Vitis vinifera* in the presence of dried flowers of *Madhuca indica* in potable drinking water. Dried flowers of *Woodfordia fruticosa* were added after the water volume lowered to one-quarter of the initial volume for initiation of fermentation while using Jaggery as a source of sugar. After fermentation was completed in about 45 days, the formulation was duly filtered and kept in sterile dark glass bottles after passing through bacterial filters of 0.2-micron pore size for further use. The preparation method was modified from the Ayurvedic Pharmacopoeia of India following Sharangadhar Samhita and Vaisajyaratnavali references for different “Arishta” formulations. All plant parts were procured from the local market and identified by qualified Ayurveda practitioners with formal academic degrees for practicing Ayurveda [16–18].

Loperamide hydrochloride (Imodium brand of Johnson and Johnson Consumer Inc.) was used as a standard drug against diarrhea induced by the effects of castor oil.

Castor oil (Erand Tail brand of Dabur) was used to induce diarrhea in animals.

Ultrapure Kaolin from Himedia Laboratories was used as a marker for determining the residual effect of AP-01 on the gastric motility of animals.

### Experimental animals

36 Sprague Dawley rats of either sex, aged around six weeks, weighing 80 to 120 grams, were obtained from the animal house of the Department of Zoology, University of North Bengal. Animals were acclimatized for seven days in standard environmental conditions, and a pellet diet of chow and water was provided in copious amounts. Experiments were conducted after the animals were kept deprived of both food and water for 18 to 20 hours. The study was in accordance with the National Research Council’s Guide for the care and use of Laboratory Animals and approved by the Institutional Animal Ethics Committee of the University of North Bengal, Siliguri, Distr. Darjeeling, West Bengal, India (Ref. No. IAEC/NBU/2018/04 dated 12.09.2018)[19].

### Microorganisms with pathogenesis and non-pathogenic properties

Different human pathogenic bacterial clinical isolates were thankfully received from the Gastro-intestinal Tract Pathogens Repository or GTPR of the National Institute of Cholera and Enteric Diseases, sponsored by the Indian Council of Medical Research (ICMR), the apex body to conduct clinical and pre-clinical research in India. Based on availability at the repository and varying degrees of resistance against multiple antibiotics, four different strains were chosen for performing in vitro tests with AP-01. All four strains were received in sealed vials with extensive care and caution and further experiments were conducted aseptically in Bio-safety level II laboratory (Omics Laboratory) as per the guidance of ICMR. Proper disposal by autoclaving all media containing GTPR strains and thereafter incinerating the contents of the autoclave were followed. *Escherichia coli 01241^2^* resistant to Ceftazidime, Aztreonam, Ceftriaxone, Trimethoprim/Sulphamethoxazole, Nalidixic Acid, Ampicillin and Cefotaxim, *Salmonella typhi 01396* enterica serovar resistant to Streptomycin, Amoxyclav, Ciprofloxacin, Norfloxacin, Doxycycline, and Erythromycin, *Salmonella typhi 01263* enterica serovar resistant to Ofloxacin, Ciprofloxacin, Nalidixic Acid and Norfloxacin along with *Shigella boydii 01399* strain resistant to Nalidixic Acid, Ciprofloxacin, Cefotaxim and Kanamycin were received from GTPR[20]. All four GTPR strains, along with a laboratory isolate of a non-pathogenic nature, *Escherichia coli K12*, were treated with polyherbal formulation AP-01 to conduct further research after determining the susceptibility/resistance pattern of the microorganisms.

Antibiotic resistance/susceptibility profiles of the above-mentioned bacterial strains were also determined at the lab as per the guidelines of CLSI (Clinical Laboratory Standards Institute) (CLSI, 2013) using the Kirby-Bauer disk-diffusion method. The laboratory findings mostly corroborated with the resistance details available with all GTPR strains. The zone of inhibition was measured and compared with breakpoints given by EUCAST/CLSI to determine the resistance/susceptibility profile of each strain against antibiotics belonging to different classes [21[22].

### Phytochemical Screening of the formulation

Primary phytochemical studies were done following standard biochemical qualitative and analytical assay procedures and literature surveys for individual plant material and the polyherbal formulation to confirm the presence of phytoconstituents [25]. Finally, the polyherbal formulation was characterized by different spectroscopic techniques like FTIR spectroscopy (Nicolet 6700 FT-IR spectrometer, Thermo Fisher Scientific, USA), 1HNMR spectroscopy (600 MHz, Bruker) and high-performance liquid chromatography-tandem mass spectrometry (LC-MS/MS, Waters, USA).

### Antimicrobial Assay of polyherbal AP-01 against GTPR^3^ pathogenic strains

The Minimum Inhibitory Concentration (MIC) of polyherbal preparation AP-01 against the four GTPR strains and *Escherichia coli K12* was determined by serial broth dilution method by adhering to the Clinical Laboratory Standards Institute (CLSI, 2013) reference method[21]. Polyherbal preparation AP-01 was added to sterile 5ml Muller Hinton broth in different concentrations from 10 µl/ml to 200 µl/ml with a gradual increment of 5 µl/ml and then 1% aliquot of the mid-log-phase culture of 0.5 McFarland standard of each bacterium was inoculated in every tube equally and incubated for 24 h at 37^0^C. After incubation, growth was measured in a spectrophotometer (SPECTROSTAR Nano - BMG LABTECH) at 600nm to determine the minimum inhibitory concentration (MIC). All the experiments were done in triplicates and repeated twice. Overnight grown mid-log phase bacterial culture was inoculated both in the presence of polyherbal AP-01 at MIC dose and without the presence of polyherbal AP-01 at 37^0^C and incubated for six hours to compare the optical density-based growth in both conditions using the spectrophotometer. At the same time, serially diluted aliquots from cultures grown under both conditions (with and without adding AP-01) were grown over Luria agar plates through the spread plate technique for reassurance of growth/inhibition of growth by counting the cfu of the strains. The highest dilution spread over the Luria agar plates, which did not show any growth, was considered to be the Minimum Bactericidal Concentration (MBC). In the lab, *Escherichia coli K12* was isolated and grown in Mueller Hinton Broth (Himedia) with and without AP-01. This experiment was conducted to study the inhibitory pattern and determine the MIC and MBC value of AP-01 against the strains.

Further, the modified Mueller Hinton Agar well diffusion method was also performed to determine the zone of inhibition produced by different volumes of undiluted AP-01 against each bacterial strain compared with negative control (DMSO ^4^) and positive control (Meropenem 10 μg). 100 μl aliquot of mid-log-phase culture of 0.5 McFarland standard containing 1 x 10^9^ cfu/ml of each bacterium was aseptically spread over 20 ml Mueller Hinton Agar plates, and sterile 6mm well borers were used to dig wells. Serially diluted volumes of 150 µl, 125 µl, 100 µl, 75 µl, 50 µl, and 25 µl AP-01 in DMSO (volume adjusted to 150 μl), and 150 μl DMSO were poured into individual wells, and Meropenem discs were placed in separate plates for each strain were incubated at 37° C for 24 hours to determine the zones of inhibition. Tests were performed in triplicates and presented as Mean +/- Standard deviation.

### Anti-swimming motility assay against GTPR pathogens

A sub-MIC dosage of AP-01 was added to semi-solid agar media to check possible alterations in the swimming motility phenomenon shown by any of the Salmonella strains of GTPR [26–27].

### Primary Acute Oral Toxicity Tests

After acclimatization, animals were distributed into several groups to check acute toxicity imparted by AP-01 when fed through the oral route. Standard dichromate oxidation method was applied to determine the level of ethanol in AP-01formulationand it was determined to be present at a quantity less than 3 % v/v at the time of study (Data not given). A dosage of up to 50 ml/Kg for the body weight of rats was found to be safe. Further studies were carried out using 5ml/Kg and 10ml/Kg dosing of AP-01 as rationalized in some earlier studies using plant extracts [28]8-29] and according to the Organization of Economic Corporation Development Guidelines [30] guidelines following the principles and criteria summarized in the Humane Endpoints Guidance Document. The rationale for such dosing was based on the body surface area ratio for extrapolating human dose to animals.

### Cytotoxicity Study

The human intestinal epithelial cell line IEC 18 and rat intestinal epithelial cell line RIE -1 were respectively procured from the National Centre for Cell Science, Pune, and Centre for Cell and Molecular Biology, Hyderabad, India, for performing MTT assay with different strengths of AP-01 and by diluting AP-01 with DMSO. Dulbecco’s Modified Eagle Medium, fortified with 0.25% Sodium bicarbonate solution and 8% fetal bovine serum, was used during the maintenance of the cell culture. CO2 incubator (Thermo Fischer Scientific). The cell lines were stored at 37°C in a 5% CO2 atmosphere with 90% humidity. An equal number of cells were placed at 96 well plates with 800 μl culture media. After 24 hours of incubation, the media was replaced by 18.75 μl, 37.5 μl, 75 μl, 150 μl, and 750 μl AP-01 after adjusting the volume to 800 μl by adding DMSO and another well with 800 μl DMSO as control. After incubation under a similar storage condition for 24 hours, the entire contents of each well were replaced by 50 μl of 5mg/ml MTT solution freshly prepared in phosphate buffer saline and incubated for another 6 hours. Succinate dehydrogenase enzyme present in cell lysates of viable cells would therefore be reacting with yellow MTT solution (3-(4,5-dimethylthiazol-2-yl)-2,5-diphenyl-2H-tetrazolium bromide) to form violet-blue crystalline Formazan molecule that could be dissolved in propan-3-ol or isopropyl alcohol for visible spectroscopy for quantification. As metabolically active cells could only produce Formazan crystals, their quantification was used to determine the number of living cells. The maximum amount of Formazan should be formed in the well containing DMSO as control. The plates were horizontally shaken after adding 100 μl of isopropyl alcohol for 2-3 mins. Absorbance was taken at λ_max_ of 600 nm, concerning the violet-blue color of Formazan at 490 nm, by 96 well plate reader (Clario Star plus, BMG Labtech). The percentage of cytotoxicity was calculated by comparing the OD value of AP-01 treated cells with the control. (OD_Control_ – OD_Treatment_)/ OD_Control_ x 100.

AP-01 was added to the cell lines as 0.25 - MIC, 0.5 – MIC, MIC, 2X MIC, and 10X MIC dosage towards lab isolate bacterial strain E. coli K12 [31–32].

### Experimental Design for the action of AP-01 on the symptom of diarrhea induced by ricinoleic acid from Castor Oil

The animals were divided into six groups, each group containing six randomly selected but comprising equal numbers of male and female Sprague Dawley rats for a parallel experimental design to study the effect of AP-01 on Castor oil-induced diarrhea. The normal group received only distilled water (10ml/Kg p.o.), and the negative control group received only Castor oil at a dose of 10ml/Kg. The positive control group was fed with a per oral dose of 2mg/Kg Loperamide hydrochloride suspended in distilled water before half an hour after receiving Castor oil (10ml/Kg *per oral*). The first two test groups received 5ml/Kg and 10ml/Kg AP-01 consecutively before half an hour of receiving Castor oil (10ml/Kg) through the oral route. However, the last test group received AP-01 at a dose of 10ml/Kg after half an hour of feeding Castor oil at a dose mentioned previously [28–29]. Each animal was individually observed under a glass funnel lined with blotting paper until twenty-four hours after feeding Castor oil and water in the case of the standard group. Observations were done to test the parameters like the onset of symptoms of diarrhea employing excreting wet feces, cessations of diarrhea by means of the end time for the expulsion of wet feces, the total weight of feces, weight of wet feces, total number of stool pellets and the total number of wet stool pellets. The percentage of inhibition of symptoms of diarrhea was determined by comparing the number of wet stools expelled during the observation period by animals of the positive control group and test groups compared to the number of wet stools removed by negative control group animals.

### The residual effect of AP on body weight, food, and water intake, and gastric motility (using Kaolin as a marker) before and after intervention

Each animal was weighed one day before performing the experiment and again weighed after 18-20 hours from the conclusion of the observation time of the investigation. Food and water intake were calculated one day before intervention by providing a fixed amount of pellet diet and a fixed volume of water and calculating the left-over mass of food and volume of water after twenty-four hours. Dietary intake and water intake were again measured after twenty-four hours from the interim time of twelve hours of observation of the experiment, as all animals were fed with a fixed amount of diet and water after twelve hours of observation.

Before the feeding of animals after the observations, each animal was administered with 2ml/Kg *per oral*. Identified from previous studies, a dose of 40% aqueous solution of Kaolin could be used as a marker of gastric motility rate as it could alter the color of stool pellets without affecting the drug action administered previously. Therefore, the measured time gap between the feeding of Kaolin and expulsion of fecal matter marked with Kaolin for each animal could serve as a determinant of gastric motility because it could be assessed as the time taken by the Kaolin to get a passage through the gastro-intestinal tract of the rats. Longer latency before the expulsion of Kaolin-marked fecal matter could be suggestive of decreasing gastric motility rates[29][33].

### Statistical analysis

The observational and all measured values were reported as Mean ± Standard Error of Mean (S.E.M.). Percentage changes compared to normal values and control group data were carried out as and when required. Paired and unpaired “students” “t” test was employed for statistical analysis for validation of inference at a level of significance of P<0.05 was considered as statistically significant. Results for antimicrobial activity tests were interpreted as Mean +/- standard deviation.

## Results

### Presence of probable phytochemicals

The presence of different functional groups viz., hydroxyl (alcoholic and phenolic-OH), carbonyl (C=O), alkoxy(C-O), epoxy (C-O-C), C-H, C=C, and C-N bonds was confirmed by the Fourier Transform Infrared (FT-IR) spectra of our polyherbal formulation taken in liquid form (**Figure 1A**)[34]. The high intense peak at 3380 cm-1 confirms the predominant presence of hydroxyl groups (alcoholic/phenolic) in the present formulation. This also indicates the considerable existence of alcohols and polyphenols in our formulation, responsible for its significant pharmacological activity. Besides, the 1H-NMR spectra of our current polyherbal formulation further indicate the presence of primarily aliphatic protons (δ;1.25 - 4.9ppm) and alcoholic/phenolic/benzylic protons (chemical shift values, δ 4.26, 4.57, and 4.9 ppm (**Figure 1B**).

**Fig. 1.**
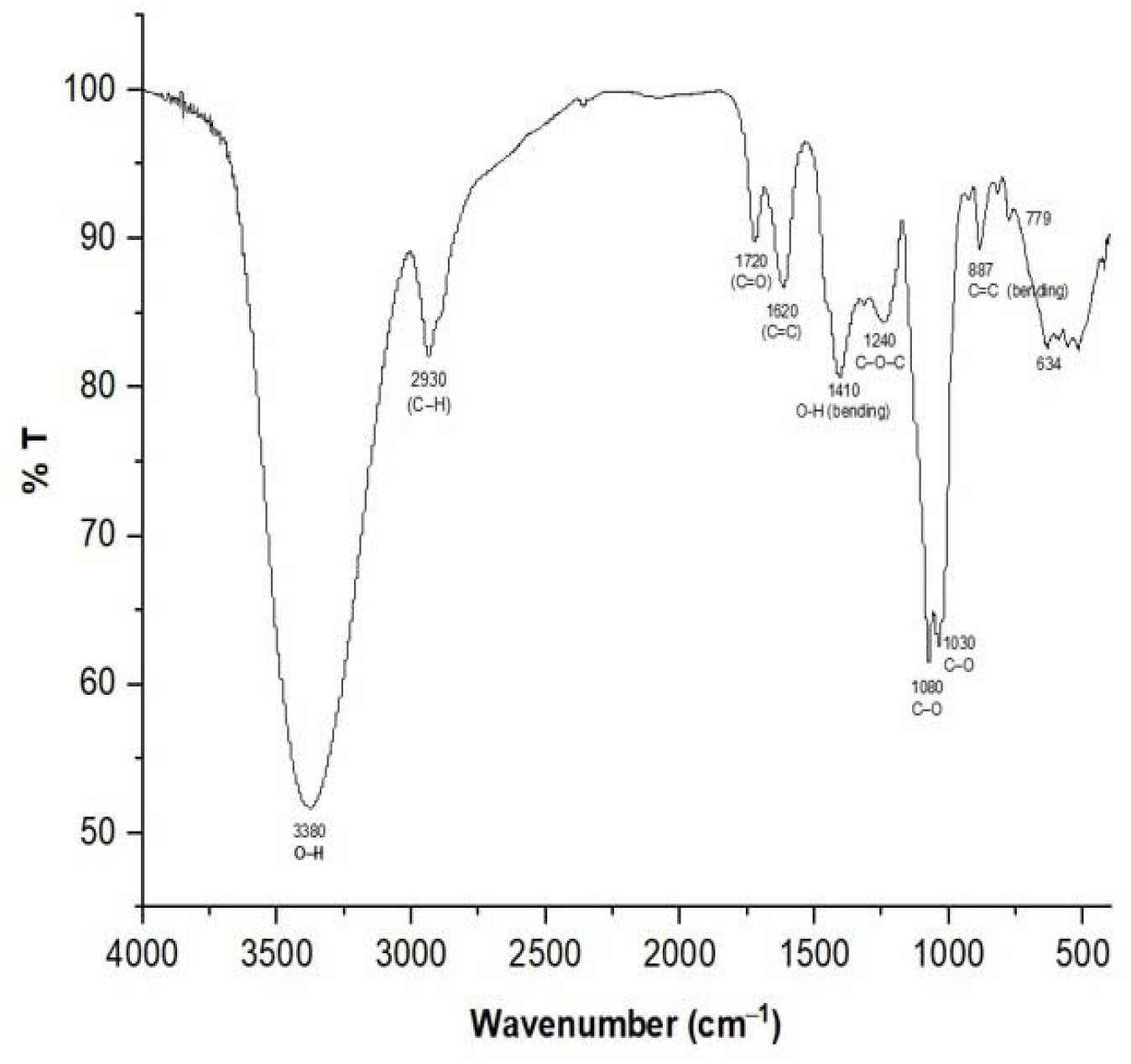
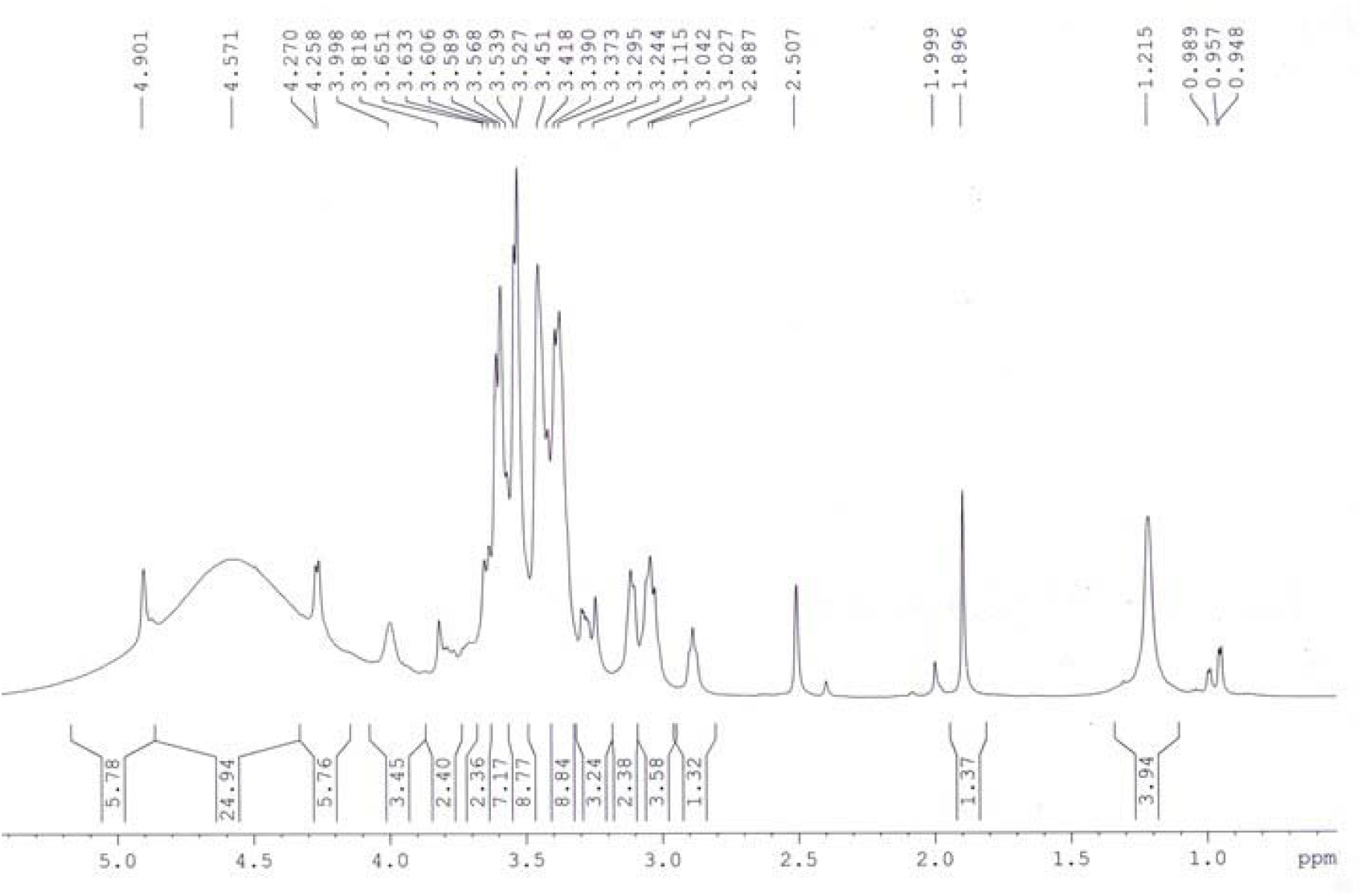
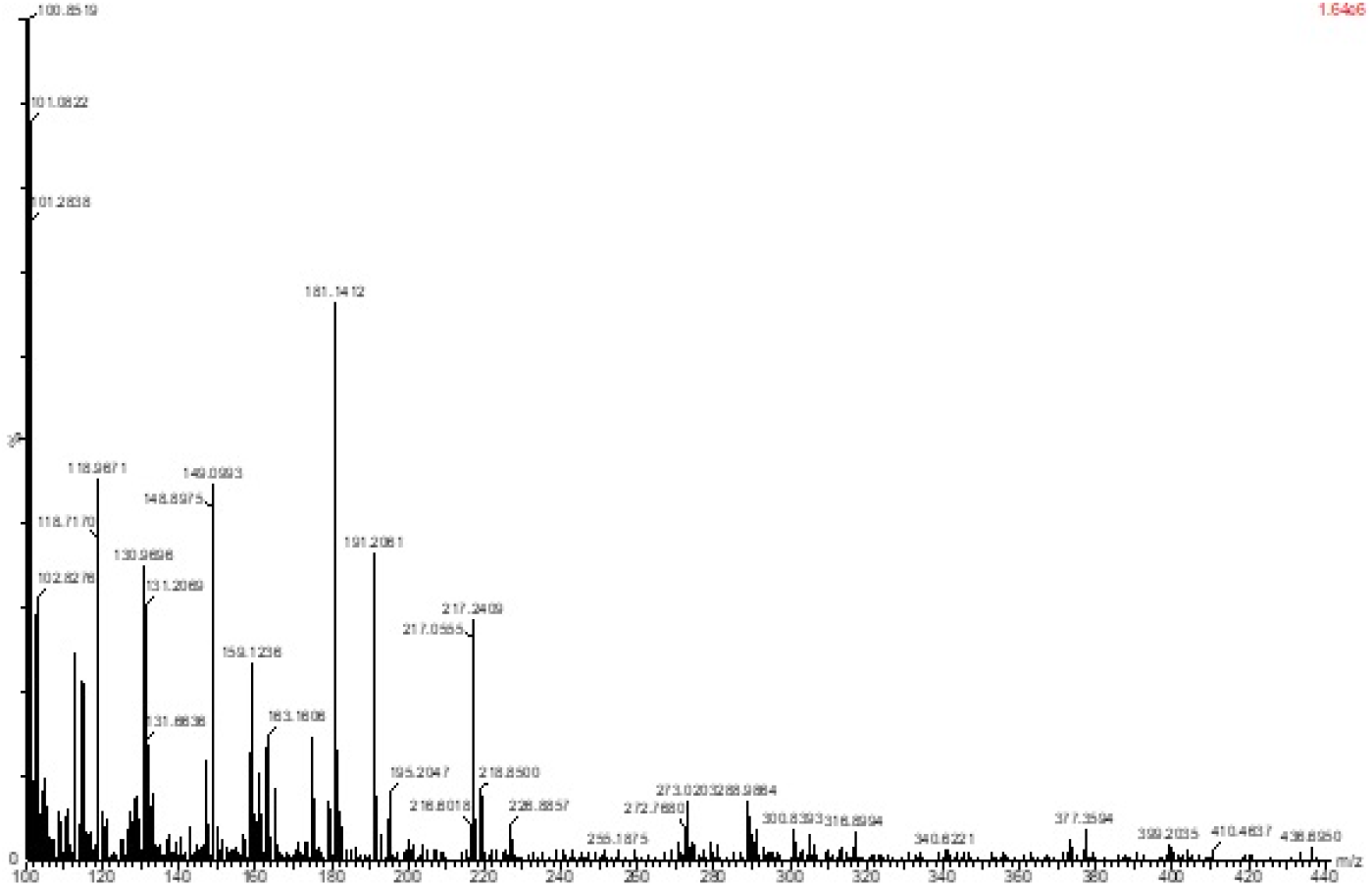
(A) FTIR; (B) NMR and (C) LC-MS study of fermented polyherbal formulation AP-01.

On account of the FT-IR and 1HNMR spectra along with the LC-MS/MS data (**Figure 1C**), the presence of diverse phytochemicals viz., quercetin (302.23), β-sitosterol (M.W. 414.7), n-octacosanol (MW-395.72), kurchine (MW-342.6), conessine (MW 356.58), antidysentericine (MW-356.5), conimine (MW-328.53), gmelinol (MW-402.4), leuteolin (MW-286.24) gmelanone (368.33), sigmasterol (MW-412.69), betulinic acid (MW-456.700), myricetin (MW-318.235), catechin (290.2681), kaempferol (MW-286.236), beta-Amyrin (MW-426.717), luteolin (MW-286.23), n-hexacosanol (MW-382.70), oleanic acid (MW-456.7), linolenic acid (MW-278.42), mannitol ( MW-182.17), myricetin (MW-318.23) and their various glycosidic derivatives is primarily inferred as bioactive compounds. The chemical composition was also compared with the literature on individual plant extracts. Hence, the major classes of phytochemicals present in our polyherbal formulation are alkaloids, flavonoids, and polyphenols.

### Cytotoxicity analysis

AP-01 was not found to be producing any cytotoxic effect in either concentration as 0.25 - MIC, 0.5 – MIC, MIC, 2X MIC, and 10X MIC dosages to either human or rat cell lines as at the highest concentration; cell viability was not found to be reduced for more than 10 percent as represented in **Table 1**.

**Table 1.**
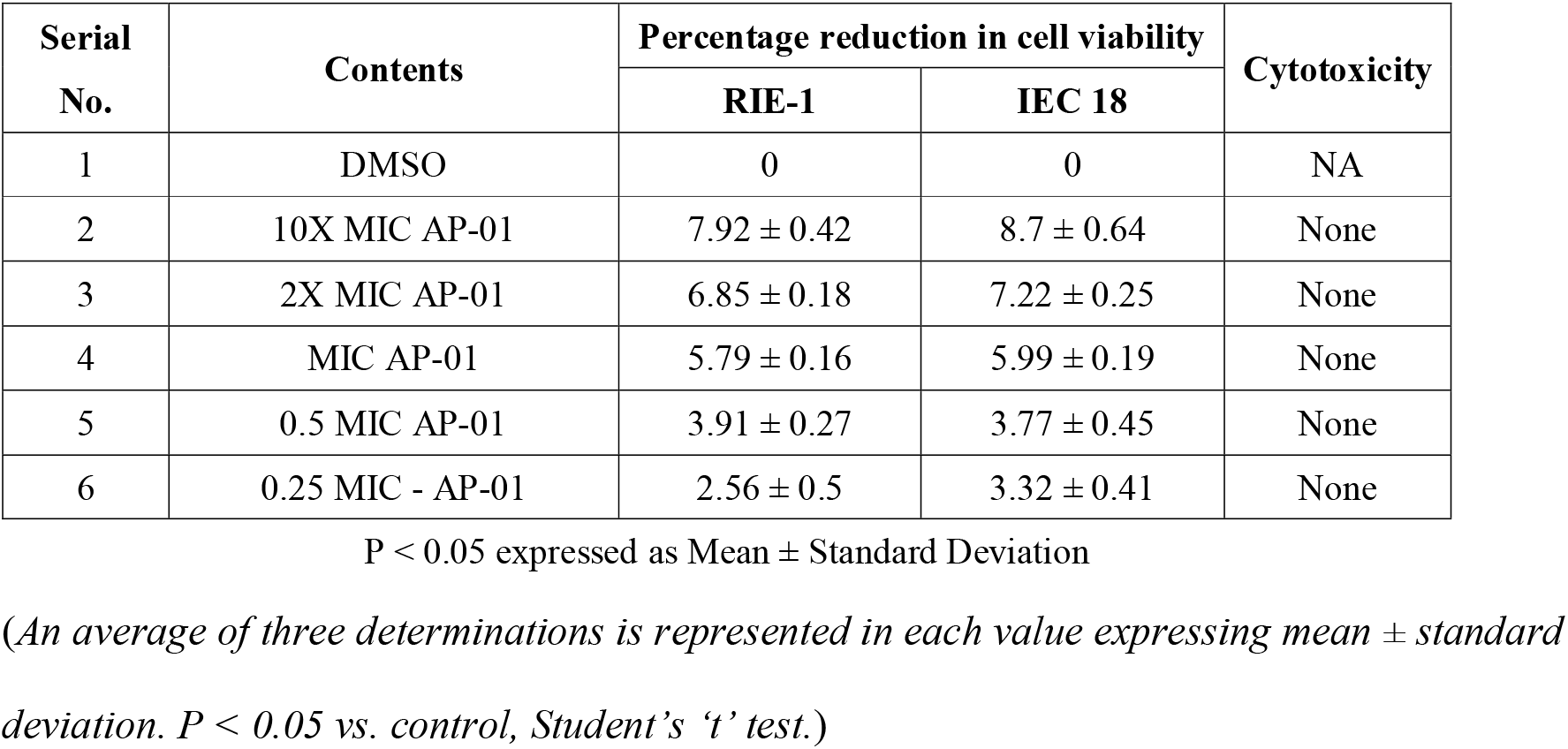
MTT Assay based Cytotoxicity study of fermented polyherbal formulation.

| Serial No. | Contents | Percentage reduction in cell viability |  | Cytotoxicity |
| --- | --- | --- | --- | --- |
|  |  | RIE-1 | IEC 18 |  |
| 1 | DMSO | 0 | 0 | NA |
| 2 | 10X MIC AP-01 | 7.92 ± 0.42 | 8.7 ± 0.64 | None |
| 3 | 2X MIC AP-01 | 6.85 ± 0.18 | 7.22 ± 0.25 | None |
| 4 | MIC AP-01 | 5.79 ± 0.16 | 5.99 ± 0.19 | None |
| 5 | 0.5 MIC AP-01 | 3.91 ± 0.27 | 3.77 ± 0.45 | None |
| 6 | 0.25 MIC - AP-01 | 2.56 ± 0.5 | 3.32 ± 0.41 | None |

P < 0.05 expressed as Mean ± Standard Deviation
(An average of three determinations is represented in each value expressing mean ± standard deviation. P < 0.05 vs. control, Student's 't' test.)

Data are presented as mean ± SD from two independent experiments, each run-in triplicate. The % inhibition was calculated as the percent difference between growth in DMSO and growth in the presence of AP-01 after 24 h of incubation.

### Effect of AP-01 on animals with diarrhea induced by Castor Oil

All fecal excrements were collected, weighed, and analyzed for the study (**Figure 2**). During the period of observation for twenty-four hours after dosing with Castor oil, all animals of the negative control group showed symptoms of diarrhea with an early onset of the release of a wet fecal pellet within an hour (an average of 52 minutes) and the symptoms persisted as long as nineteen hours. Animals belonging to the first and second test groups showed dose-dependent delayed onset of symptoms; however, the occurrence ceased at a significantly earlier time (three hours) in the case of rats pre-treated with 10ml/kg AP-01 formulation. The third test group, which consisted of animals that received AP-01 after half an hour of Castor oil feeding, also showed a delayed onset of symptoms compared to the negative control and first test groups. Treatment with Loperamide as a standard drug produced a significant reduction of symptoms at a rate of 83.1%, while test groups having 5ml/Kg and 10ml/Kg AP-01 dosages as pre-treatment and 10ml/Kg AP-01 as post-treatment with Castor oil consecutively reduced occurrence of diarrhea at a percentage inhibition of 40.11%, 79.71% and 60.84% as shown in **Table 2**. Duration of diarrhea was found to be significantly decreased in the case of the positive control group and second test group (that received 10ml/Kg AP-01 pre-treatment) and measured consecutively as 94 minutes and 140 minutes. The normal animal group did not show any symptoms of diarrhea throughout the experiment and observational period.

**Fig. 2.**
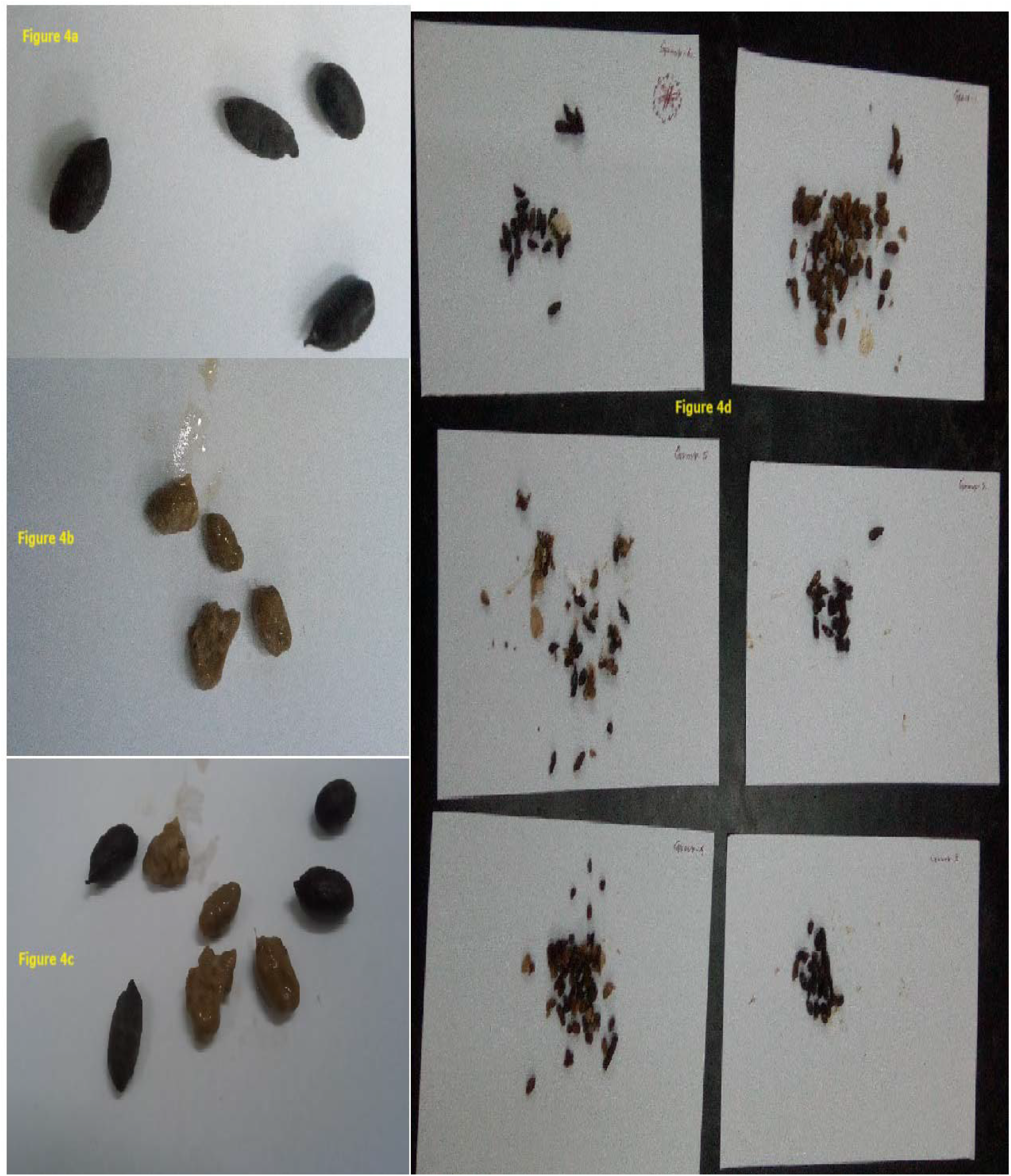
(Colour) Comparative normal and wet fecal materials of all animal groups.

**Table 2.** Effect of AP-01 on castor oil (0.8 ml) induced diarrhoea in Sprague Dawley rats.

| Group | Onset of Diarrhoea (Mins) | Cessation of Diarrhoea (Mins) | Total wt. of stool (g) | Total wt. of wet stool (g) | Total no. of stool | Total no. of wet stool | % of inhibition |
| --- | --- | --- | --- | --- | --- | --- | --- |
| Normal (Water) | None | None | 0.1385±0.001 | 0 | 3.66±0.21 | 0 | N.A. |
| -ve control (Castor oil) | 51.83±0.54 | 1185.66±29.32 | 1.8433±0.07 | 1.7366±0.06 | 37.83±0.56 | 35.33±0.71 | 0% |
| +ve control (C.oil+Loper.) | 214.66±4.76 | 308±6.63 | 0.1868±0.007 | 0.1768±0.007 | 6.83±0.4 | 6±0.365 | 83.01% |
| 5ml/kg AP (before C.oil) | 101.83±1.78 | 602.33±6.56 | 0.9318±0.007 | 0.9146±0.008 | 23.16±0.6 | 21.16±0.7 | 40.11% |
| 10ml/kg AP (before C.oil) | 180.66±3.18 | 320.66±5.05 | 0.1911±0.005 | 0.1795±0.005 | 8.33±0.49 | 7.16±0.48 | 79.71% |
| 10ml/kg AP (After C.oil) | 91.66±1.23 | 395.5±8.04 | 0.472±0.003 | 0.4596±0.002 | 15.66±0.42 | 13.83±0.31 | 60.84% |

An average of six determinations is represented in each value expressing mean ± standard error of mean.
P < 0.05 vs. control, Student's 't' test.

### The residual effect of AP-01 on body weight, food, and water intake, and gastric motility

Expulsions of stool pellets marked with Kaolin were visually altered in color (**Figure 3**). Kaolin-marked stool pellets took a significantly longer time to pass through the gastrointestinal tract of animals in the case of both the groups treated with 10ml/Kg AP-01, however, a significantly higher latency period was shown when the dose of AP-01 was given at a later period (in the third test group). Body weight and food intake were reduced significantly in the case of the negative control group receiving only Castor oil; however, there were no significant changes in the case of body weight or food intake for the animals belonging to any test groups and positive control group. There was a significant increase in food intake for the animals of the normal group. Another significant change was shown in **Table 3** depicting the increase in water intake only for the animals belonging to the negative control group.

**Fig. 3.**
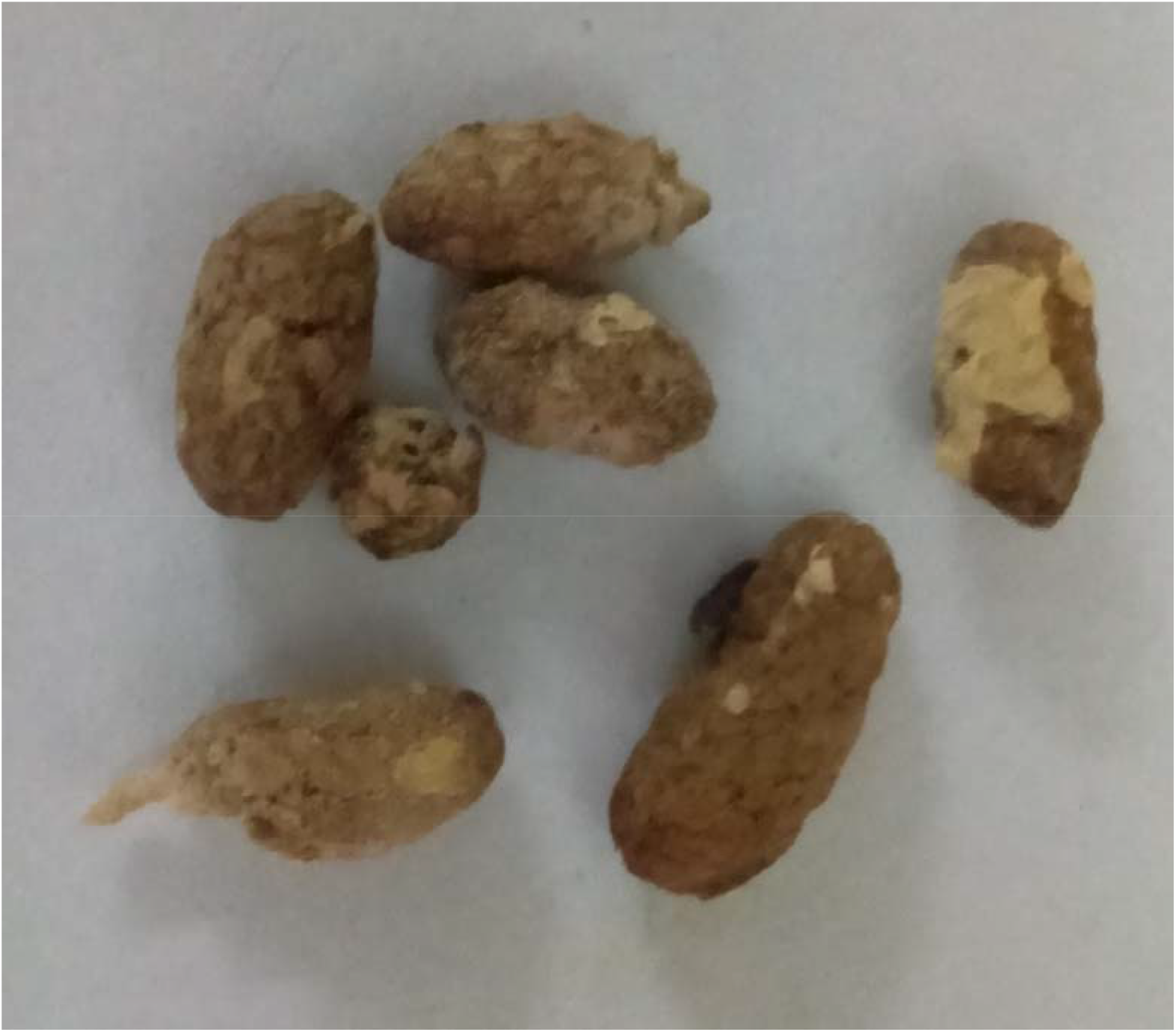
(Colour) Kaolin expulsion through faeces as a marker of gastric motility.

**Table 3.** Residual effect of AP-01 on body weight, food and water intake and gastric motility of Sprague Dawley rats.

| Group | Body Weight(g)<br>(pre-treat) | Body Weight(g)<br>(post-treat) | Food intake<br>(pre-treat)<br>(g) | Food intake<br>(post-treat)<br>(g) | Kaolin<br>expulsion rate | Water<br>intake(ml)<br>(pre-treat) | Water intake(ml)<br>(post-treat) |
| --- | --- | --- | --- | --- | --- | --- | --- |
| Normal<br>(Water) | 106±5.65 | 107.83±5.16 | 15.95±0.824 | 16.48±0.814 | 367.67±0.882 | 16.5±0.730 | 16.58±0.889 |
| -ve control<br>(Castor oil) | 107.5±4.66 | 102±4.99 | 16.17±0.688 | 15.18±0.714 | 326.67±2.551 | 15.33±0.654 | 17.5±0.592 |
| +ve control<br>(C.oil+Loper.) | 98.16±5.96 | 99.5±6.32 | 14.75±0.87 | 14.93±0.956 | 377.67±2.403 | 15.42±0.898 | 14.67±0.802 |
| 5ml/kg AP<br>(before C.oil) | 95.66±5.41 | 96.66±5.29 | 14.43±0.818 | 14.52±0.783 | 371.17±1.797 | 14.42±0.8 | 14.17±0.76 |
| 10ml/kg AP<br>(before C.oil) | 99.16±5.47 | 101.33±5.48 | 14.93±0.754 | 15.02±0.878 | 397.5±1.91 | 15.17±0.882 | 14.75±0.642 |
| 10ml/kg AP<br>(After C.oil) | 102.83±5.42 | 103.16±5.14 | 15.38±0.787 | 15.48±0.79 | 409±4.412 | 15.67±0.872 | 15.5±0.774 |

*An average of six determinations is represented in each value expressing mean ± standard error of mean.*
*P < 0.05 vs. control, Student's 't' test.*

### The Effect of AP-01 on Gastric Pathogens Collected from Gastro-intestinal Tract Pathogens Repository

*E. coli 01241*, *S. typhi 01263*, *S. typhi 01396* & *Shigella boydii 1399* strains of pathogenic bacteria were collected from NICED to conduct in vitro antimicrobial tests on human pathogenic bacteria with AP-01. Primary tests confirmed the inhibitory effect. AP-01 was found efficacious against NICED-GTPR Multiple Antibiotics Resistant human pathogenic strain *E. coli 01241* (Resistant to CAZ, AT, CTR, SXT, NA, AMP, CTX); *S. typhi 01396* (Resistant to S, AMC, CIP, NOR, DO, E); *S. typhi 01263* (Resistant to OFX, CIP, NA, NOR) and *Shigella boydii 1399* (Resistant to NA, CIP, CTX, K). (**Table 4**). The minimum inhibitory concentrations of AP-01 against each GTPR strain, determined through the broth dilution method and well diffusion method was 80 μl/ml v/v *E. coli 01241* and 100 μl/ml v/v for the other three strains. MIC dose of AP-01 for the lab isolate *E. coli K12* was determined to be 75 μl/ml v/v (**Tables 5 and 6**).

**Table 4.** Antibiotics resistance patterns based on CLSI/EUCAST zone of Inhibition breakpoints.

| Serial No. | Antibiotic Agent | Symbol | Disc Content | CLSI Breakpoint (Resistance) mm or less | Eucast Breakpoint (Resistance) mm or less | Experimental Data *E. coli* 01241 (mm) | Experimental Data *S.typhi* 01396 (mm) | Experimental Data *S.typhi* 01263 (mm) | Experimental Data *S.boydii* 01399 (mm) |
| --- | --- | --- | --- | --- | --- | --- | --- | --- | --- |
| 1 | Amikacin | AK | 30 mcg | 14 | 13 | 17 | 15 | 17 | 19 |
| 2 | Amoxyclav | AMC | 30 mcg | 13 | 19 | 21 | **11 [R]** | **18 [R]** | **13 [R]** |
| 3 | Ampicillin | AMP | 10 mcg | 13 | 14 | **11 [R]** | 15 | 19 | **12 [R]** |
| 4 | Aztreonam | AT | 30 mcg | 17 | 21 | **14 [R]** | 22 | 23 | 22 |
| 5 | Ceftazidime | CAZ | 30 mcg | 19 | 17 | **12 [R]** | 21 | 22 | 24 |
| 6 | Ceftriaxone | CTR | 30 mcg | 19 | 20 | **11 [R]** | 22 | 24 | 23 |
| 7 | Ciprofloxacin | CIP | 5 mcg | 20 | 20 | 23 | **13 [R]** | **18 [R]** | **13 [R]** |
| 8 | Cefotaxime | CTX | 30 mcg | 22 | NA | **12 [R]** | 23 | 26 | **14 [R]** |
| 9 | Doxycycline | DO | 30 mcg | 10 | 10 | 13 | **9 [R]** | 12 | 17 |
| 10 | Erythromycin | E | 15 mcg | 13 | 16 | 18 | **10 [R]** | 19 | 20 |
| 11 | Kanamycin | K | 30 mcg | 13 | NA | 16 | 24 | 17 | **11 [R]** |
| 12 | Nalidixic Acid | NA | 30 mcg | 13 | 12 | **11 [R]** | 17 | **10 [R]** | **10 [R]** |
| 13 | Norfloxacin | NX | 10 mcg | 12 | 12 | 19 | **9 [R]** | **9 [R]** | **9 [R]** |
| 14 | Ofloxacin | OF | 5 mcg | NA | 19 | 21 | **15 [R]** | **17 [R]** | 22 |
| 15 | Streptomycin | S | 10 mcg | 11 | NA | 14 | **8 [R]** | 19 | 17 |
| 16 | Trimethoprim/<br>Sulphamethoxazole | SXT | 5 mcg | 10 | 15 | **8 [R]** | 16 | 17 | 16 |
| 17 | Cefepime | CPM | 30 mcg | 18 | 21 | **13 [R]** | 23 | 22 | 20 |
| 18 | Doripenem | DOR | 10 mcg | 19 | 21 | 26 | 25 | 21 | 22 |
| 19 | Imipenem | IPM | 10 mcg | 19 | 16 | 24 | 23 | 23 | 21 |
| 20 | Meropenem | MRP | 10 mcg | 19 | 16 | 23 | 23 | 22 | 22 |

[R] Designates resistance towards this antibiotic agent

**Table 5.** MIC and MBC of polyherbal AP-01 against bacterial strains of study.

| Serial No. | Bacterial Strains | MIC $\mu\text{l/ml}$ | MBC $\mu\text{l/ml}$ |
| --- | --- | --- | --- |
| 1 | *Escherichia coli K12* | 75 | 90 |
| 2 | *Escherichia coli 01241* | 80 | 120 |
| 3 | *Salmonella typhi 01396* | 100 | 150 |
| 4 | *Salmonella typhi 01263* | 100 | 150 |
| 5 | *Shigella boydii 01399* | 100 | 100 |

**Table 6.** Zone of inhibition shown by different volumes of AP-01 while keeping DMSO as negative control and Meropenem as positive control.

| Bacterial Strains | Meropenem | AP01 Undiluted | AP01 in DMSO (volume adjusted to 150 $\mu\text{l}$) | | | | | DMSO |
| --- | --- | --- | --- | --- | --- | --- | --- | --- |
| | 10 mg | 150 $\mu\text{l}$ | 125 $\mu\text{l}$ | 100 $\mu\text{l}$ | 75 $\mu\text{l}$ | 50 $\mu\text{l}$ | 25 $\mu\text{l}$ | 150 $\mu\text{l}$ |
| *Escherichia coli K12* | 25.67 ± 0.58 | 25 ± 0.5 | 24.83 ± 0.58 | 25 ± 0.87 | 24.83 ± 0.29 | 16.75 ± 0.35 | 12.5 ± 0.5 | 0 |
| *Escherichia coli 01241* | 23.83 ± 0.29 | 22.67 ± 0.29 | 22.5 ± 0.5 | 22.33 ± 0.29 | 22.16 ± 0.29 | 15.67 ± 1.15 | 11.5 ± 0.5 | 0 |
| *Salmonella typhi 01396* | 23.67 ± 0.29 | 22.83 ± 0.58 | 22.67 ± 0.58 | 22.5 ± 0.5 | 16.17 ± 0.29 | 14.17 ± 0.76 | 13 ± 0.87 | 0 |
| *Salmonella typhi 01263* | 22.83 ± 0.76 | 22.33 ± 0.29 | 22.33 ± 0.29 | 21.83 ± 0.76 | 15.33 ± 0.58 | 15.17 ± 1.04 | 13.33 ± 0.58 | 0 |
| *Shigella boydii 01399* | 22.67 ± 0.58 | 21.17 ± 0.29 | 20.17 ± 0.76 | 20.17 ± 0.29 | 17.83 ± 0.76 | 15.67 ± 0.76 | 14.17 ± 0.29 | 0 |

P < 0.05 expressed as Mean ± Standard Deviation

### Anti-swimming motility assay against GTPR pathogens

The presence of a sub-MIC dose of AP-01 (50 μl/ml v/v) in semi-solid media significantly arrested the swimming motility of GTPR strain *S. typhi 01263* which was found to be the only strain to produce a swimming motility phenomenon. In the following figure (**Figure 4**), the first left side figure showed the swimming motility observed without the presence of AP-01 in the media, and the adjoining figure showed the inhibition of swimming motility in the presence of sub-MIC dose of AP-01 in media.

**Fig. 4.**
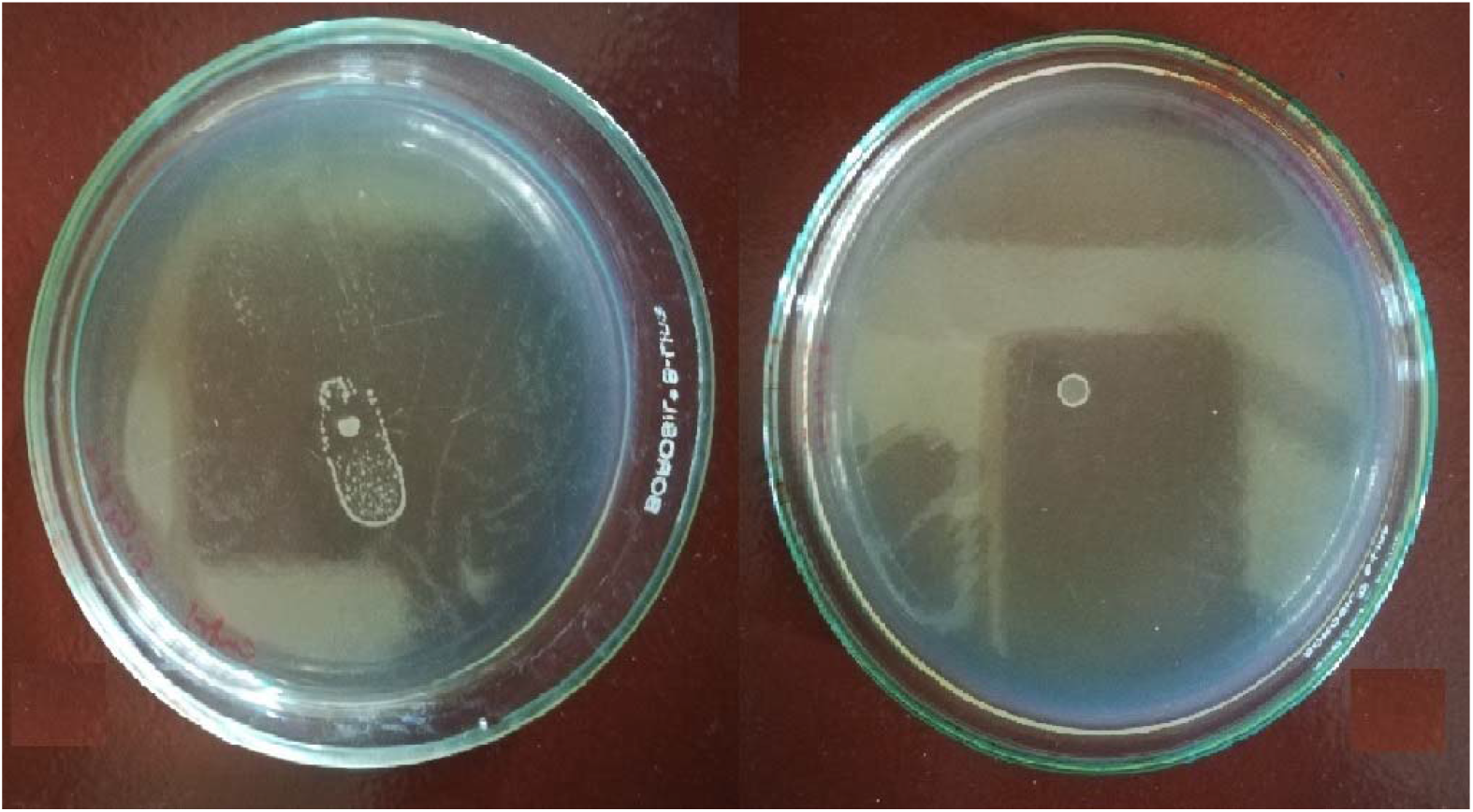
(Colour) Anti swimming motility effect shown by AP-01.

## Discussion

Castor oil induces symptoms of diarrhea by releasing ricinoleic acid through the action of lipases in the gastric mucosa of mammalian animals. Ricinoleic acid triggers the release of prostaglandins following inflammatory activity produced on the upper gastrointestinal tract. Alteration in permeability of gastric mucosal cells occurs with the release of prostaglandins, and a decrease in absorption of electrolytes like Na+ and K+ happens as a result. Due to enhanced permeability and the hypersecretory effect of gastric mucosa, peristalsis hastens, and the symptoms of secretory diarrhea appear [2–3]. Formulations using *Holarrhena antidysenterica* stem bark as a major constituent was previously reported to be effective against secretory diarrhea. From the results of the current study, Ayurveda-based fermented polyherbal formulation AP-01 was found to be significantly effective against diarrhea induced by Castor oil, suggesting that the polyherbal formulation could produce its action through anti-secretory mechanisms of action. The reduction of diarrhea in terms of the number of wet tools in the case of test groups was significant and comparable to the treatment effects of a standard drug used in positive control groups. The reduction of symptoms of diarrhea was found to be dose-dependent; however, there was a valuable finding that AP-01 could exert its anti-secretory action even after the introduction of the diarrhea-causing substance. AP-01 not only delayed the onset of symptoms when administered prior but also reduced the time of incidence of appearance of symptoms and finally ceased any effects of Castor oil-induced diarrhea.

Regularization of gastric motility signifies the restoration of normalcy after treatment with anti-diarrheal and anti-secretory agents. The longer duration of action of such drugs helps in increasing dose interval and stimulating patient compliance. The recent study had put some light on the anti-motility effect of the polyherbal formulation AP-01. The assessment of in vivo passage of the drug was indirectly done by using a marker (Kaolin) which lightens the color of stool which can be easily distinguished from the normal or wet stool color of rats[33], without affecting the pharmacodynamics. Significant hindrance of gastric motility was observed with AP-01 in comparison to standard drugs where no significant effects were noticed after six to seven hours of administration. The efficacy of AP-01 in reducing gastric motility and frequency of defecation was found to be most significant when an equal dose of AP-01 was administered one hour after in the case of test group three in comparison to test group two, suggesting some residual anti-motility and anti-secretory effect pertained by AP-01 as it might still be active on gastric mucosa in prior to metabolism, excretion, and degradation of the formulation. The significant decrease in body weight and food intake in the case of the negative control group suggested that the symptoms of diarrhea disappeared, and the appetite and gastric condition were yet to be recovered even after more than one day after treatment whereas the animals of other group were apparently returned to normalcy. Further proof of this hypothesis was ascertained from the results showing a significant increase in water uptake in the case of a negative control group, whereas other groups showed no significant change. It could possibly be caused by excessive loss of water and electrolytes from the gastric mucosa of the animals fed with only castor oil, leaving them with a higher demand of water for the body to quench the loss. Animals belonging to all other groups were shown no significant change in water intake as osmolarity and water balance in the gastrointestinal tract and in the whole body respectively had become close to normal. Reduction of motility without significantly altering amounts of food or water intake after treatment was shown with the administration of AP-01 at different dosages and administration time proved the formulation to be efficacious yet safe in case of secretory diarrhea.

The Ayurveda-based “Arishtha” preparatory method was applied in formulating AP-01 that suggested boiling for a definite time would not affect the efficacy of the active constituents exerting pharmacological actions; therefore provided us with further scope in modification of dosage form and formulation and further detailed study of chemical and biochemical constituents that could lead us in demonstrating the exact mode of action of this polyherbal formulation, prepared by following Ayurveda in treating diarrhea and dysentery of various etiology. However, in vitro studies using individual plant components or unfermented polyherbal mixture did not show any promising results when compared with the actions exerted by the fermented polyherbal formulation AP-01 (Data not shown).

Using human pathogenic bacteria for producing diarrhea in a murine model *per oral* route of administration was deemed unsuitable and unethical due to low body weight and possible consequences leading to animal sacrifices[30]. A zoological scale-up using either a canine model or rabbit model should have to be adopted for future studies to develop a standard infectious diarrhea animal disease model using pathogenic bacteria, and it was out of the scope of the current study. Therefore, in vitro bacterial susceptibility studies were conducted with AP-01 against all four GTPR strains and the result showed significant inhibition of the pathogenic bacteria with MICs within a feasible dosing regimen. Inhibition of swimming motility of Salmonella sp. by AP-01 at a sub-lethal dose was suggestive of the formulation to be capable of hindering the wider spread of the virulent bacterial species within the affected epithelial mucosa, consequently minimizing the virulence and infection with time[35].

## Conclusion

AP-01 was definitely found efficacious against multiple antibiotic-resistant pathogens and safe for gastric mucosa, leaving an impression to research with the formulation with greater depth and detail. The mechanism of action could only be determined with further future studies including detailed chemical analysis and structure-activity relationship determination so that this unused arsenal given by nature can be modified through cutting-edge technology in developing smarter dosage forms like soft gel capsules, chewable pellets, or anything better suitable for better patient compliance and broader action spectrum.

## Authors’ contribution

Subhanil Chakraborty: Conceptualization, Data curation, Formal analysis, Investigation, Methodology, Writing – original draft. Babli Roy: Data curation, Formal Chemical Analysis, Investigation, Writing – original draft. Subhajit Sen: Data curation, Formal analysis. Santi M. Mandal: Data curation, Formal analysis, Investigation. Ranadhir Chakraborty: Conceptualization, Data curation, Funding acquisition, Investigation, Methodology, Project administration, Supervision, Writing – review & editing.

## ACKNOWLEDGEMENT

The authors are also thankful to Dr. Min Bahadur, Professor, and Dr. Tilak Saha, Assistant Professor, Department of Zoology, University of North Bengal, for their active cooperation in the procurement of animals and necessary animal ethical permissions from the Department of Zoology, University of North Bengal. Author S.C. is grateful to Dr. Anoop Kumar, Assistant Professor, Department of Biotechnology, University of North Bengal, and to late Dr. Prof. Dhananjoy Pal, Bengal School of Technology for their kind support in performing the cytotoxicity study. The authors are grateful to Doctor Subrata Roy, B.A.M.S. of J.B.Roy State Ayurvedic Medical College and Hospital for the identification of plant parts for preparing polyherbal formulation AP-01 and for providing valuable suggestions. The authors are very thankful to Dr. Ranjan Kumar Nandy and Dr. Asish Kumar Mukhopadhyay of NICED, Kolkata, sponsored by ICMR for supplying pathogenic strains from their Gastrointestinal Tract Pathogen Repository (ICMR Sanction No. 5/8-1(231)/2008-ECD-II) without any cost.

## Footnotes

2 Identification number assigned to each clinical isolate by GTPR

3 GTPR signifies Gastric Tract Pathogen Repository of Indian Council of Medical Research, Govt. of India

4 DMSO is Dimethyl sulphoxide deviation.

